# Cryo-EM structure of TMEM63C suggests it functions as a monomer

**DOI:** 10.1101/2023.07.04.547736

**Authors:** Yuqi Qin, Daqi Yu, Dan Wu, Jiangqing Dong, William Thomas Li, Chang Ye, Kai Chit Cheung, Yingyi Zhang, Yun Xu, YongQiang Wang, Yun Stone Shi, Shangyu Dang

## Abstract

The TMEM63 family proteins (A, B, and C), calcium-permeable channels in animals that are preferentially activated by hypo-osmolality, have been implicated in various physiological functions. Deficiency of these channels would cause many diseases including hearing loss. However, their structures and physiological roles are not yet well understood. In this study, we determined the cryo-EM structure of the mouse TMEM63C at 3.56 Å, and revealed structural differences compared to its plant orthologues OSCAs. Further structural guided mutagenesis and electrophysiological studies demonstrated the important roles of the coupling of TM0 and TM6 in channel activity. Additionally, we confirmed that the physiological state of TMEM63C is monomer, while TMEM63B is a mix of monomer and dimer in cells, suggesting that oligomerization is a novel regulatory mechanism for TMEM63 proteins.

## Introduction

The response of cells to both internal and external mechanical stimulation is critical for living organisms to adapt to and survive in their environments. The mechanosensitive ion channels (MSCs) are responsible for cellular mechanotransduction, a process that converts mechanical force to electrical and ion flux signals upon receiving mechanical stimulations including physical force and osmotic pressure^1,2^. Several types of MSCs have been identified^3^, including TREK/TRAAK two-pore domain potassium channels (K2P)^4^, Piezo1/2 non-selective cation channels^5^, NompC/TRPN from transient receptor potential (TRP) superfamily^6^, epithelial sodium channel/degenerin (ENaC/DEG) superfamily sodium channel^7^, TMC1/2^8^, and TMEM63/OSCA^9^.

TMEM63s, together with its plant orthologues OSCAs, are members of a newly identified OSCA/TMEM63 cation channel family that are activated by mechanical forces of osmolality^9^. OSCAs are characterized as calcium-permeable channels sensing hyperosmotic stress and influencing water transpiration regulation and root growth in plants^10^, and several structures about OSCA are available^11–14^. Different from OSCAs, TMEM63s calcium influx is preferentially activated by hypo-osmolality in animals^15^, and none of the TMEM63 structures were determined. Though remotely related in protein sequences to members of transmembrane protein 16 (TMEM16), the OSCA/TMEM63 family proteins share similar 3-dimensional (3D) architectures with TMEM16s^16–18^. Furthermore, TMC^19^ also resembles structures of TMEM16s. Based on the structural similarities, TMC, OSCA/TMEM63 and TMEM16 are grouped into the TMEM16 superfamily with diverse functions^3^.

TMEM63 has three members, TMEM63A, B and C in most animals including mouse and human. In Drosophila, only one type of TMEM63 is identified that senses humidity^20^ and food grittiness^21^. Mouse TMEM63s, and human TMEM63A were reported to induce stretch-activated currents when expressed in naïve cells^9^. According to clinical and genome-wide association studies, heterozygous missense mutations of human TMEM63A (G168E, I462N, and G567S) are linked to an infantile disorder, showing similar symptoms resembling hypomyelination leukodystrophy^22^. In support, cells transfected with these disease-related variants in TMEM63A did not generate stretch-activated currents^22^. TMEM63B plays an important role in maintaining auditory function in mouse outer hair cells (OHCs)^15^. In addition, overexpressed TMEM63B in HEK293T cells can enhance cell migration and wound healing^23^. The increased level of TMEM63B mRNA expression detected in ductal carcinoma T47-D human tumor cell line suggests that TMEM63B may also serve as cancer biomarker^23^. TMEM63C is essential for maintaining the kidney filtration barrier integrity in zebrafish, and expresses low in chronic kidney disease patients^24^.

Despite these clinical and experimental observations, studies of TMEM63s are hampered by the lack of structure information and physiological roles. Here, we determined the 3.56-Å structure of mouse TMEM63C by using single-particle cryo-electron microscopy (cryo-EM), which worked as a monomer in cells. This cryo-EM structure, in combination with structure-guided mutagenesis and electrophysiological studies, allows us to establish the structural mechanisms and the physiological functions of TMEM63C.

## Results

### Overall Structure of TMEM63C

The full-length mouse TMEM63C was overexpressed in HEK293S cells. Target protein was solubilized and purified in the detergent n-Dodecyl-β-d-maltoside (DDM), followed by exchange into lauryl maltose neopentyl glycol (LMNG) for structural studies (Supplementary Figure 1a). Using single-particle cryo-EM, we determined the structure of TMEM63C at 3.56 Å resolution as a monomer (Figure 1a-b, Supplementary Figure 1b-d, Supplementary Figure 2-3, Supplementary Table 1). Local resolution analysis revealed a higher resolution in the transmembrane regions and a lower resolution in the intracellular regions, suggesting the dynamic of intracellular regions (Supplementary Figure 3a). TMEM63C contains 11 transmembrane helices, TM0-TM10. The N-terminus is oriented towards the extracellular, while the C-terminus faces the intracellular (Figure 1c). Between TM2 and TM3, there is a long cytosolic domain mainly formed by α-helices (Figure 1c). TM3 and TM4 are tilted to the membrane, and situated at the outer boundary of the transmembrane region (Figure 1a-b). A short α-helix is linked to the intracellular region of TM6 and TM7 separately. TM7 and TM8 are shorter that do not extend the whole membrane (Figure 1).

**Figure 1.**
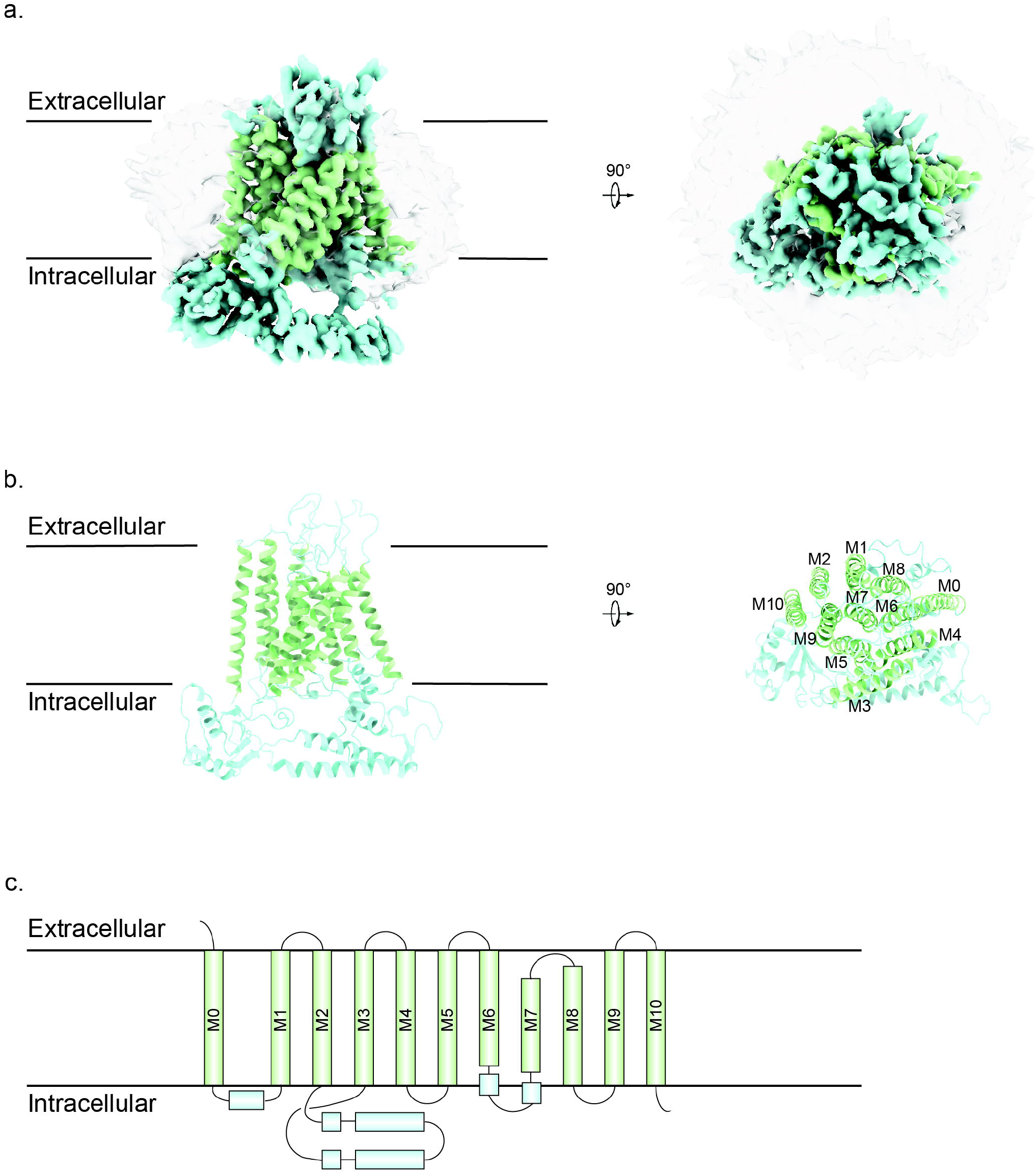
Cryo-EM structure of TMEM63C. (**a**) Side view (left) and top view (right) of TMEM63C EM density map at 3.56 Å resolution. The transmembrane region and soluble region are colored in green and blue respectively. The apparent micelle displays at a lower threshold. (**b**) Side view (left) and top view (right) of TMEM63C model. The color scheme is the same as the EM density map. (**c**) Topology of TMEM63C. The transmembrane helices are labeled M0 to M10.

**Figure 2.**
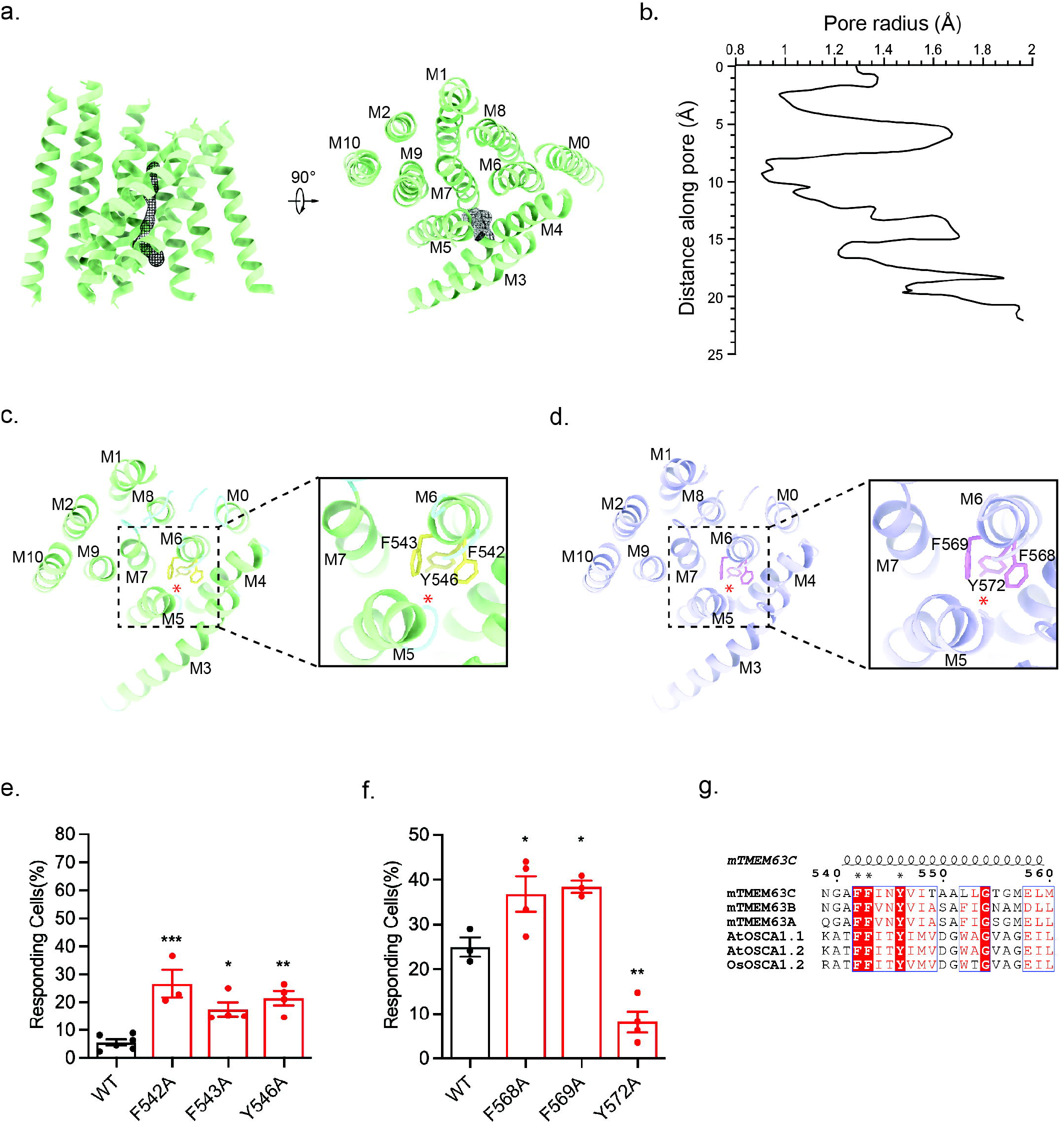
The pore profile of TMEM63C. (**a**) Top view (left) and side view (right) of TMEM63C transmembrane α-helices (green) and putative pore (black mesh). The pore is surrounded by α-helices M3 to M7 predicted by caver. (**b**) The pore radius along distance of TMEM63C. (**c**) The three bulky pore-facing residues F542, F543 and Y546 (yellow) of TMEM63C in the pore. (**d**) The three bulky pore-facing residues F568, F569 and Y572 (pink) of TMEM63B in the pore. The position of the pore is indicated with a red asterisk. The TMEM63B protein structure (purple) is predicted using Swiss-Model based on the TMEM63C structure. (**e**) The responding ratio TMEM63C expressing cells to hypo-osmolality stress. The point mutation F542A, F543A and Y546A exhibit a higher responding ratio compared with the wild type TMEM63C. (**f**) The responding ratio of TMEM63B expressing cells to hypo-osmolality stress. The point mutation F568A and F569A exhibit a higher while the Y572 exhibit a lower responding ratio compared with the wild type TMEM63B. (**g**) Sequence alignment of TMEM63 and OSCA about the M6 transmembrane α-helix (coil). The three highly conserved bulky pore-facing residues F, F and Y are indicated with black star.

**Figure 3.**
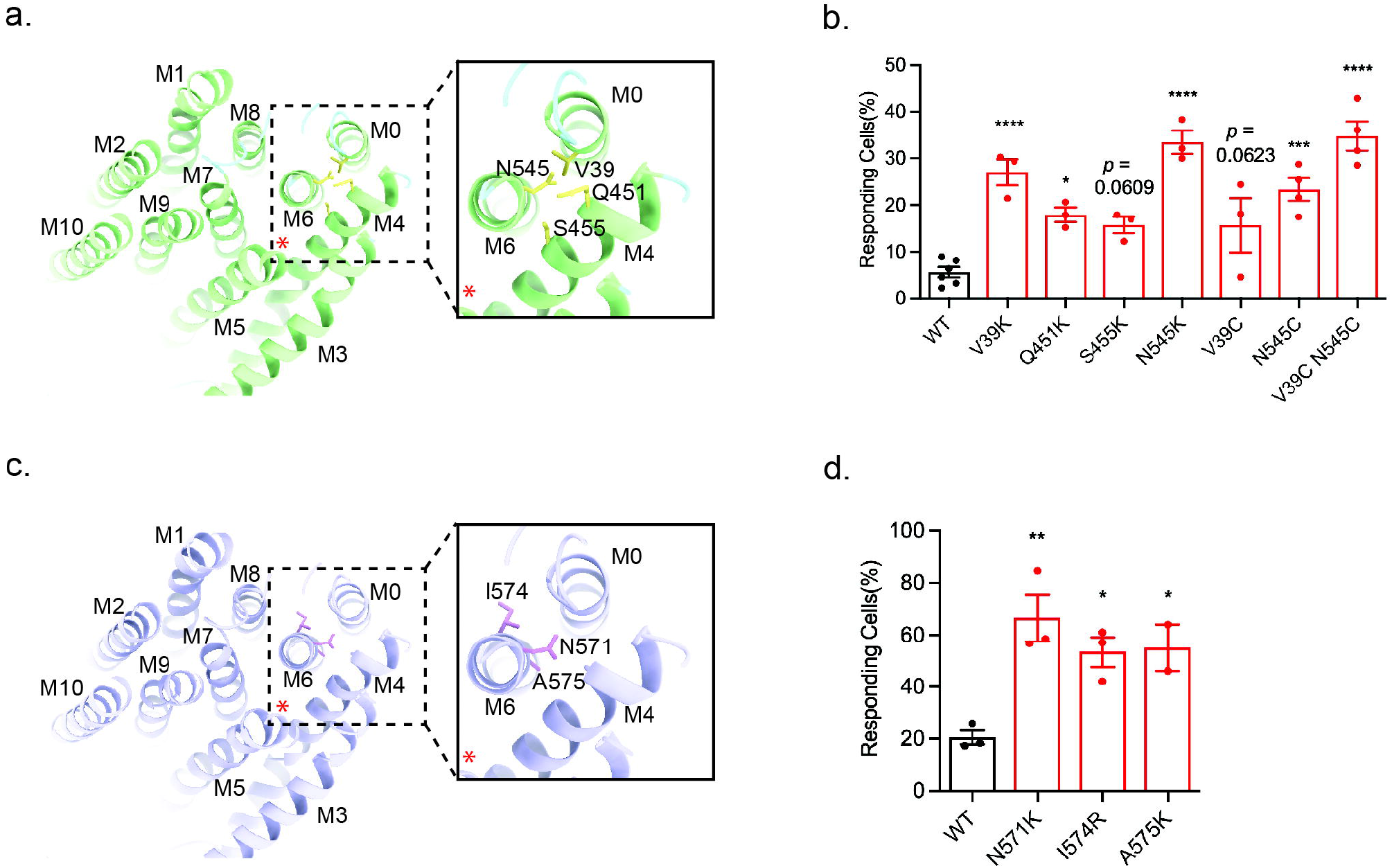
Coupling of TM0 and TM6 is important for channel activity. (**a**) The TMEM63C residues V39 in M0, Q451, S455 in M4 and N545 in M6 (yellow) outside the pore. The position of the pore is indicated with red star. (**b**) The responding ratio TMEM63C expressing cells to hypo-osmolality stress. The point mutation V39K, Q451K, S455K, N545K, V39C, N545C and V39C N545C exhibit a higher responding ratio compared with the wild type TMEM63C. (**c**) The TMEM63B residues N571, I574 and A575 in M6 (pink) outside the pore. The position of the pore is indicated with a red asterisk. (**d**) The responding ratio of TMEM63B expressing cells to hypo-osmolality stress. The point mutation N571K, I574R and A575K exhibit a higher responding ratio compared with the wild type TMEM63B.

Despite the monomeric TMEM63C observed in the cryo-EM map, the overall structure of TMEM63C resembles OSCA, its plant orthologue. mTMEM63C and atOSCA1.1 (PDB:6JPF)^14^ exhibit a sequence identity of 21.6% for the full length (Supplementary Figure 4), while transmembrane region RMSD is 1.25 for pruned pairs and 3.77 for all pairs. TMEM63C and OSCA consist of 11 transmembrane α-helices spanning from TM0 to TM10 with conserved structures. Additionally, the intracellular soluble domain of both proteins is primarily composed of the region between TM2 and TM3, as well as the C-terminus.

**Figure 4.**
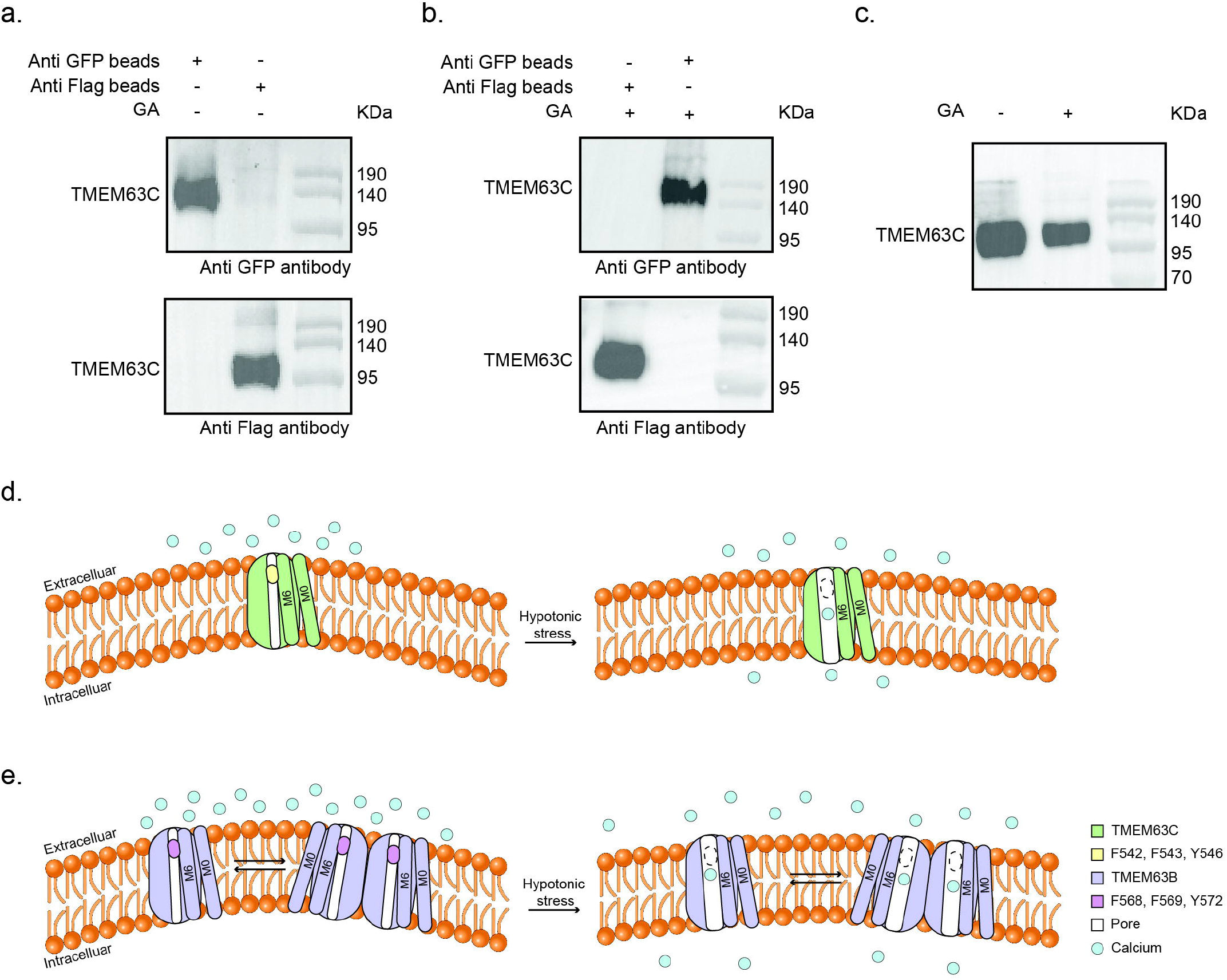
Native state of TMEM63C is a monomer. (**a**) Co-IP of TMEM63C without crosslinking. (**b**) Co-IP of TMEM63C with crosslinking before extraction by using detergent. (**c**) Crosslink of TMEM63C-flag in membrane with GA. (**d-e**) Proposed working model for TMEM63C (d) and TMEM63B (e). Calcium ion is indicated as blue circle.

Besides a similar fold, some differences still exist in the pore region of TMEM63C and OSCA. Particularly, the short α-helix located at the cytoplasmic side of TM6 swung to TM4 to avoid the clash with TM0 (Supplementary Figure 5a-b), the intracellular half of which moved close to TM6, compared to that in OSCA. Other pore-forming helices, including the TM5, TM7, also shifted slightly, thereby contributing to a different pore profile of TMEM63C (Supplementary Figure 5c).

Using pore-forming helices as reference, structural alignment of TMEM63C and OSCA indicated the major differences come from helices surrounding the pore region (Supplementary Figure 5). The extracellular side of the TM2 adopted a shift to TM10. Consequently, two neighboring helices, TM9 and TM10, were pushed away rigidly. On the other hand, the cytoplasmic side of TM7 slightly shifted to TM1, and pushed it away from the pore region. The conformational differences of these pore-region surrounding helices may sense membrane distortion introduced by mechanical forces, and convert it to intracellular signal by regulating channel activity. In OSCA, TM2 and TM10 face the dimer interface and connect to the dimer interaction intracellular domain. These morphological differences in TM2 and TM10 might also contribute to oligomerization state variation between TMEM63C and OSCA.

### Pore profile of TMEM63C

Similar to OSCA, the ion-passing pore of TMEM63C is surrounded by TM3-7. The Caver program^25^ was used to measure the radii of the pore region. The results show that TMEM63C is in closed state with two restriction sites, the radii of which is smaller than 1 Å (Figure 2a-b). Three bulky pore-facing residues (F542, F543 and Y546 on TM6) (Figure 2c, Supplementary Figure 6a), highly conserved among TMEM63 and OSCA (Figure 2g), have been proposed to play critical roles in channel activities^14^. To test this hypothesis, we examined the channel activity of TMEM63C by combining functional assays and mutagenesis. Expectedly, alanine substitution of F542, F543 and Y546 all resulted in an increased percentage of responding cells compared to wild type TMEM63C (Figure 2e). These results further demonstrated three highly conserved bulky residues play important roles in channel gating. Under normal conditions, three bulky hydrophobic resides, localized at the entry site of the ion passing pore, prevent access of the hydrophilic calcium ions. Once hypotonic condition is achieved, these residues may point away from ion passing pore as a consequence of TM6 rotation.

Interestingly, lysine substitution of V39 on TM0, Q451 and S455 on TM4, and N545 on TM6 all elevated the ratio of responding cells in TMEM63C (Figure 3a-b, Supplementary Figure 6c). These residues are not facing pore region, the substitution of lysine, a relatively bulky positive charged residue, may either cause rotation of pore helices, like TM6, or destabilize the pore region, to avoid steric repulsion, therefore increasing the opening probability of TMEM63C.

Previously, MD simulation and mutagenesis results suggested that TM6 and TM0 may couple together to trigger the opening of the pore in OSCA upon mechanical stimuli^14^. To test if this mechanism could be applied to TMEM63C, we performed cysteine substitution of V39 on TM0 and N545 on TM6, two residues are spatially close to each other based on the structural observation. As expected, TMEM63C variants with single mutation (V39C, N545C) presented a higher activity than wild type. Moreover, double mutant (V39C and N545C) produced the highest activity (Figure 3b). These results suggested coupling of TM0 and TM6 may play important role in channel opening in responding to the hypotonic environment.

Relatively high sequence similarity (44% identity and 63% similarity) of TMEM63B and TMEM63C suggested that the structure of TMEM63B should be similar to that of TMEM63C (Supplementary Figure 4, Supplementary Figure 8). Therefore, the homology model of TMEM63B was generated based on sequence alignment and atomic model of TMEM63C. Similar to TMEM63C, we performed mutagenesis and functional assay for three highly conserved bulky residues located in the entry site of the pore. Alanine substitution of F568 and F569 on TM6 of TMEM63B led to increased channel activity, while Y572A reduced channel activity (Figure 2d, f, Supplementary Figure 6b). Substitution of residues on TM6 (N571, I574, A575) with lysine or arginine all increased the channel activity at least two times higher compared to that of wild type TMEM63B (Figure 3c-d, Supplementary Figure 6d). These results suggested although TMEM63 proteins share similar working mechanisms, there might be subtle differences to fulfill their functional varieties under physiological conditions.

### TMEM63C function as a monomer

Surprisingly, the monomeric TMEM63C was observed in the cryo-EM structure. To investigate its physiological functional state, we performed pull-down assay by attaching GFP or FLAG at the C-terminus of TMEM63C, respectively. The cells were transfected by viruses of both TMEM63C-FLAG and TMEM63C-GFP together for overexpression. Following the same protocol as TMEM63C for structural studies, the harvested cells were solubilized in DDM before loading to GFP nanobody (GFP-Nb) beads and FLAG beads, respectively. After washing off the nonspecifically bound proteins, the beads were analyzed by western blotting. The results clearly show that sample from GFP beads can only be detected by GFP antibody, whereas sample from FLAG beads can only be detected by FLAG antibody, suggesting no dimer/oligomer were formed for TMEM63C (Figure 4a). To exclude the possibility that the dimeric TMEM63C may be disassembled when extracted from the membrane, we purified cell membrane and performed crosslinking before isolating by DDM. Consistently, no TMEM63C-FLAG/TMEM63C-GFP proteins have been observed for sample binding to GFP/FLAG beads (Figure 4b). Besides, the TMEM63C-FLAG protein was expressed and crosslinked in the membrane without extraction. No dimer/oligomer band with higher molecular weight can be detected (Figure 4c). These observations further demonstrated that TMEM63C is a monomer in the cell membrane.

## Discussion

The TMEM63/OSCA family has been characterized as a mechanosensitive cation channel family in 2018^9^. The plant orthologues OSCAs have been shown to sense hyperosmotic stress as well as regulate water transpiration and root growth^9^. Several structures of OSCAs in similar closed states have been reported^11–14^. In contrast, the physiological roles of the TMEM63s remain unknown until 2020 when TMEM63B was identified as an osmosensor that is sensitive to hypotonic stress and required for hearing in mice^15^, and a humidity sensor in 2022^20^. Considering the low sequence similarities (Supplementary Figure 4, Supplementary Figure 8) and dramatically different responses to osmolality (hypo vs hyper) between TMEM63 and OSCAs, their molecular mechanisms are expected to be different.

In this study, we identified three highly conserved residues in TMEM63B/C that serve as a gate to regulate channel activity. In addition, our mutagenesis results suggested that the coupling of TM0 and TM6 played critical roles in opening the channel for ion transport. Previous studies have suggested that MSCs sense mechanical force through membrane tension and distortion. Upon stimulated by mechanical forces, membrane will be distorted and generate a thinner region surrounding MSC. Membrane thinning could cause tilting of peripheral transmembrane helices (TM0 in TMEM63 proteins), and then trigger the conformational changes of pore-forming transmembrane helices (TM6 in this case). In this study, we further demonstrated the coupling of TM0 and TM6 is important for the channel activity of TMEM63C, which could be a general mechanism for TMEM63 proteins, and even TMEM63/OSCA family proteins.

Intriguingly, different from its homology OSCAs, TMEM63C functions as a monomer, which is confirmed by both cryo-EM structure and biochemical experiments. In principle, oligomerization states should not affect the channel function of TMEM63 since each monomer contains an intact ion permeation pore. In fact, even for the structurally homologous proteins, TMEM16A^26^ and OSCAs, which form homodimers in cryo-EM structures, each monomer contains an intact ion permeation pore. We notice that several residues in the cytoplasmic region, which contribute to the dimerization of the AtOSCA1.1 by hydrogen bonding^14^, are not conserved in TMEM63 (Supplementary Figure 8), suggesting that the monomeric state of TMEM63, at least TMEM63C, may be physiological.

Although sharing high sequence similarity, TMEM63B showed dramatically higher activity than TMEM63C. Previous study suggested that TMEM63B is a dimer in physiological states^27^. The activity differences may be attributed to the oligomerization state of TMEM63B and TMEM63C, which is partially supported by our cryo-EM studies of purified TMEM63B. 2D classification of TMEM63B particles indicated that some 2D averages showed clear C2 symmetry from top/bottom views, indicating the existence of dimeric TMEM63B (Supplementary Figure 7). In contrast, 2D averages of TMEM63C only showed monomeric but not dimeric states. These results suggested differences of oligomerization in TMEM63B and TMEM63C.

On the basis of the observation in structure and channel activity, the dimerization may provide a novel strategy to regulate the channel activity of TMEM63. Although the monomeric TMEM63 contains an intact pore region and can conduct channel function independently, the activity is relatively low. Dimerization of TMEM63 may require conformational adjustment in a way that facilitates channel open probability, thus produce a higher channel activity in TMEM63B (Figure 4d, e). Future studies are needed to investigate channel activities of monomeric and dimeric TMEM63 protein to confirm this hypothesis. It would be interesting to find out whether the oligomerization states of TMEM63B will affect its mechanosensory functions and whether oligomerization is a form of regulation to fine-tune the sensitivity of ion channels.

## Materials and Methods

### Protein expression and purification

The mouse TMEM63C (Uniprot ID: Q8CBX0) was cloned into vector pEG BacMam with a C-terminus TEV cutting site and GFP tag. Bacmids were generated from DH10Bac bacteria, and baculovirus was produced and amplified in Sf9 cells according to the Bac-to-Bac protocol. The HEK293S cells were cultured in FreeStyle medium, 2% FBS, 5% CO_2_, and 40% humidity at 37 °C. 500 ml HEK293S with a density of 2.0 × 10^6^ cells/ml were infected by baculovirus to a volume ratio 1:100 (baculovirus: HEK293S). 8 hours after viral infection, 10 mM Sodium butyrate was added. HEK293S cells were collected after another 48 hours.

Cells were resuspended with buffer 50 mM Tris pH 7.0, 150 mM NaCl, 10% glycerol, and protein inhibitor cocktail, and then broken through dounce homogenizer. Membrane proteins were extracted using 1% DDM and 0.1% CHS for two hours at 4°C. The insoluble fraction was removed via centrifugation at 180,000 rpm for 45 min. The supernatant was incubated with preequilibrated GFP nanobody coupled CNBr-Activated Sepharose 4B resin for an hour at 4°C, followed by three times wash with buffer 50 mM Tris pH 7.0, 150 mM NaCl, 10% glycerol, 0.02% DDM, and 0.002% CHS. The protein was released from the resin through three hours of TEV digestion at 4°C to remove the C-terminus GFP tag. Subsequently, 500 ul concentrated protein was injected to Superdex 200 increase 10/300 column for size-exclusion chromatography with buffer 50 mM Tris pH 7.0, 150 mM NaCl, 0.002% LMNG, and 0.0004% CHS. The peak fraction was verified by SDS-PAGE and concentrated to 1 mg/ml for cryo-EM analysis.

### Cryo-EM sample preparation and data collection

4 μL purified protein was placed on the holy-carbon grids (Quantifoil Au R1.2/1.3, 300 mesh), which were glow-discharged for 30 s. The grids were blotted for 4 s with blot force 0, at 100% humidity, 4°C, and plunge-frozen into liquid ethane using Vitrobot Mark IV (Thermo Fisher Scientific).

11,058 movies were collected on 300 kV Titan Krios (Thermo Fisher Scientific) equipped with a GIF BioQuantum and K3 camera (Gatan) using software EPU (Thermo Fisher Scientific). Each movie contained 40 fractions with a total dose of ∼50 e-/Å^2^ for 4.5 s exposure time. The pixel size was 1.06 Å per pixel, and the defocus ranged from -1.0 μm to -2.5 μm.

### Cryo-EM data processing

All movies were motion-corrected, dose-weighted, and gain-normalized using MotionCor2^28^. The contrast transfer function (CTF) of non-does-weighted micrographs was estimated by Gctf^29^. 8,754,694 particles were picked by Gautomatch with 256 × 256 box size, extracted from dose-weighted micrographs in RELION^30^, and then cleaned up by using 2D classification and heterogeneous refinement in cryoSPARC^31^. The well-sorted particles generated a reference map, which worked as a reference to enrich the particles^32^ from the Gautomatch picked raw particles through seed facilitated 3D classification^33^. The raw particles were split into several portions, and each portion was combined with seed particles. These combined particles were classified using referenced and biased maps, followed by removing duplicates. The enriched 1,417,132 particles were further cleaned up by resolution gradient 3D classification^33^. To obtain the reference for the resolution gradient, a map was generated using 958,924 particles after 2D classification and seed facilitated 3D classification, then combined with the low-pass filtered 10 Å and 20 Å map. Both the seed facilitated 3D classification and resolution gradient 3D classification were done through cryoSPARC heterogeneous refinement. The 258,464 particles yielded a 3.56 Å map using local and global CTF refinement, local refinement and DeepEMhancer^34^.

### Model building

The initial model of TMEM63C was predicted with AlphaFold^35^, and fitted into cryo-EM map in Chimera^36^. The model was manually adjusted in COOT^37^ and subsequently refined with real space refinement in PHENIX^38^ iteratively. The transmembrane α-helix region of the model was identified unambiguously, while the loop region was adjusted based on the remaining density. The intracellular region (211-400) from AlphaFold prediction fits into the fragmented density. The TMEM63C model is validated by the Molprobity score^39^ and Ramachandran plots in PHENIX. Additionally, the structure of TMEM63B is predicted using homology modeling according to the TMEM63C model in Swiss-Model^40^.

### Co-IP

Two constructs TMEM63C with a C-terminus GFP tag and TMEM63C with a C-terminus Flag tag, were co-expressed in the HEK293S cells. Cells were collected and washed via centrifugation at 4,200 rpm for 10 min, followed by resuspending with lysis buffer 20 mM Hepes pH 7.0, 150 mM NaCl, 10% glycerol, and an additional protein inhibitor cocktail. 1% DDM and 0.1% CHS were added to lysis cells, and extract membrane protein at 4°C for two hours. After removing the cell pellet at 12,000 rpm for 30 min, the supernatant was divided into two parts that incubated with GFP resin and Flag resin at 4°C overnight separately. The GFP resin and Flag resin were washed at 3,000 rpm for 5 min four times. Each protein coupled resin was analyzed by SDS/PAGE and western blot using both anti-GFP and anti-Flag antibodies.

### Crosslink

The TMEM63C is expressed in the HEK293S cells, which were collected and resuspended with lysis buffer 20 mM Hepes pH 7.0, 150 mM NaCl, 10% glycerol, and an additional protein inhibitor cocktail. Cells were broken through dounce homogenizer, and the pellet was removed via centrifugation at 6,000 rpm for 30 min. From the supernatant, the membrane was obtained via ultracentrifugation at 33,500 rpm for an hour. The membrane was resuspended with lysis buffer and crosslinked with 0.02% glutaraldehyde at 4°C for 30 min. The 50 mM Tris 7.0 was applied to stop the crosslink reaction. The crosslinked membrane was analyzed by SDS/PAGE and western blot.

### SDS/PAGE and western blot

The protein samples were added to a 12% Bis-Tris gel system for electrophoresis. PVDF membranes were applied to blot the protein sample at constant 400 mA, 4°C for 2 hours, and blocked with 5% milk TBST buffer. The primary antibodies GFP tag rabbit polyAb (Proteintech) or monoclonal anti flag M2 antibody (Sigma) was incubated overnight at 4 °C, followed by three times TBST wash. The respective secondary antibodies were incubated for an hour at room temperature, followed by three times TBST wash. The signal was detected with supersignal west pico plus chemiluminescent substrate (Thermo Fisher Scientific) using ChemiDoc Imaging Systems (Bio-rad).

### Cytoplasmic Ca^2+^ measurements

The cytoplasmic Ca^2+^ influx was monitored by free calcium indicator GCaMP6f as previously reported^15,27^. TMEM63C/TMEM63B-P2A-GCaMP6f vectors were transfected into N2a cells mounted on the coverslip. Forty hours after transfection, the cells were perfused with isotonic extracellular solution containing 70 mM NaCl, 5 mM KCl, 1 mM CaCl_2_, 1 mM MgCl_2_, 10 mM HEPES, and 10 mM glucose (pH 7.4 adjusted with NaOH, 300 mOsm/liter adjusted with mannitol). The isotonic solution was exchanged to 170 mOsm/liter hypotonic solution without changing the ionic concentrations by a peristaltic pump (Longer Precision Pump, BT100-2J, China) at a constant speed. The osmolarity was measured by a vapor pressure osmometer (Wescor, Vapro 5600). The cytoplasmic calcium fluorescence was recorded at 1 Hz for 10 min by the Hamamatsu digital imaging camera (Hamamatsu, C11440-22U) at RT (24 ± 2°C) using 488 nm illumination. The change of fluorescence was normalized by the ratio of real-time intensity (Ft) relative to the initial value (F0). The cells with Ft/F0 > 1.5 were considered as positive responses to hypotonic challenge.

## Data availability

Cryo-EM density map of TMEM63C has been deposited in the Electron Microscopy Data Bank (accession no. EMD-XXXX). Atomic coordinate has been deposited in the Protein Data Bank (ID codes XXXX). All other data are available from the corresponding authors upon reasonable request.

## Acknowledgments

We thank Yifan Cheng for critical reading of the manuscript. All cryo-EM data were collected at the Biological Cryo-EM Center at HKUST, generously supported by a donation from the Lo Kwee Seong Foundation. This project is supported by grants from Hong Kong Research Grants Council (ECS26101919, GRF16103321, GRF16102822 and C6001-21EF), Southern Marine Science and Engineering Guangdong Laboratory (Guangzhou) (SMSEGL20SC01), Guangdong Basic and Applied Basic Research Foundation (2021A1515012460), Shenzhen Special Fund for Local Science and Technology Development Guided by Central Government (2021Szvup140), and HKUST start-up and initiation grants to S.D.. This work is also supported by grants from the National Key R&D Program of China (2019YFA0801603 to Y.S.S.); the National Natural Science Foundation of China (32170951 to Y.S.S., 82101393 to D.W., 81920108017, 82130036 to Y.X.); the Natural Science Foundation of Jiangsu Province (BE2019707 to Y.S.S., BK20210008 to D.W., BE2020620 to Y.X.); Jiangsu Provincial Medical Key Discipline (ZDXK202216 to Y.X.); Special Fund for Science and Technology Innovation Strategy of Guangdong Province (2021B0909050004 to Y.S.S.); the Fundamental Research Funds for the Central Universities (021414380533 to Y.S.S.); the Jiangsu Planned Projects for Postdoctoral Research Funds (2021K462C to D.W.); the STI2030-Major Projects (2022ZD0211800 to Y.X.). D.Y. and J.D. are supported by LKS fellowships.

## Author Contributions

Y.Q., J.D., W.T.L., K.C.C., and Y. W. performed protein purification and biochemical experiments. Y.Q., D.Y. J.D., Y.Z. and S.D. prepared cryo-sample and collected cryo-EM datasets, Y.Q., D.Y. and J.D. processed images, built the atomic model.

D.W. and C.Y. performed electrophysiological experiment. Y.W., Y.S.S. and S.D. supervised experiments and data analysis. Y.Q. and S.D. prepared figures and wrote the manuscript with input from all others. All authors contributed to data analysis and manuscript preparation.

