## Supplementary Figures for "Cryo-EM structure of TMEM63C suggests it functions as a monomer"

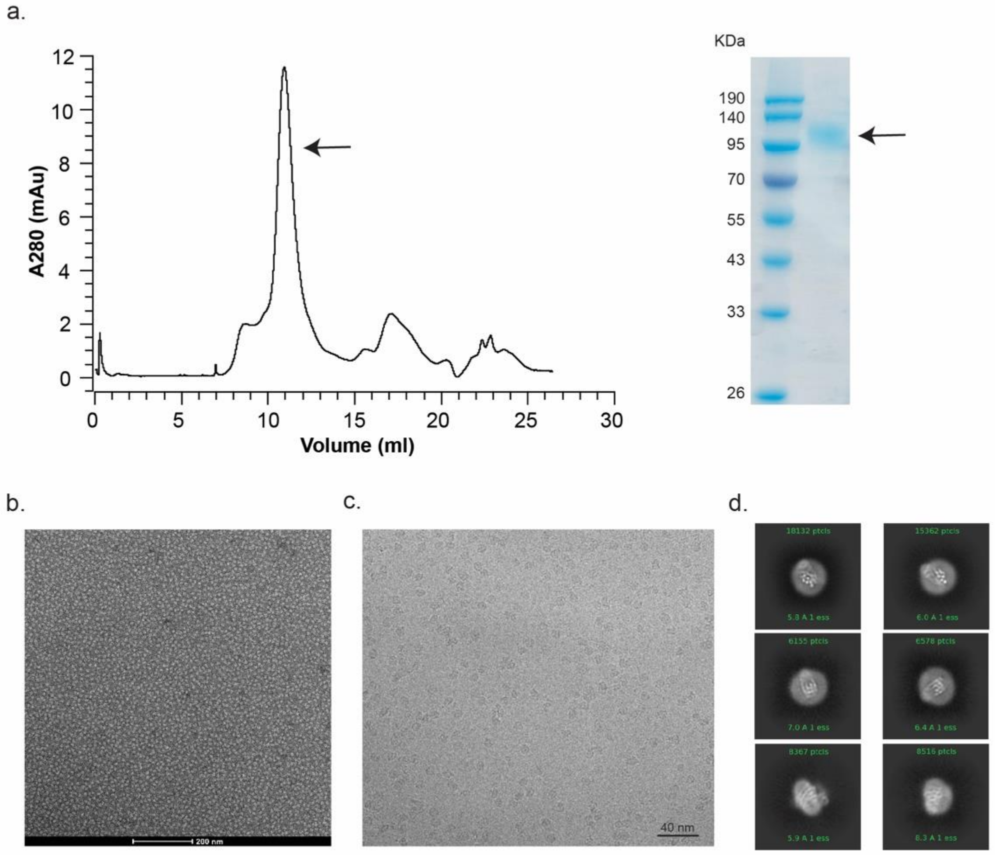


**Supplementary Figure 1 Sample preparation of TMEM63C.** (**a**) Size-exclusion chromatography of TMEM63C in detergent LMNG (left). The peak fraction is verified by Coomassie blue staining SDS-PAGE (right). (**b**) A representative negative stain micrograph of TMEM63C. (**c**). A representative cryo-EM micrograph of TMEM63C. (**d**) The 2D class average of TMEM63C final reconstituted particles.

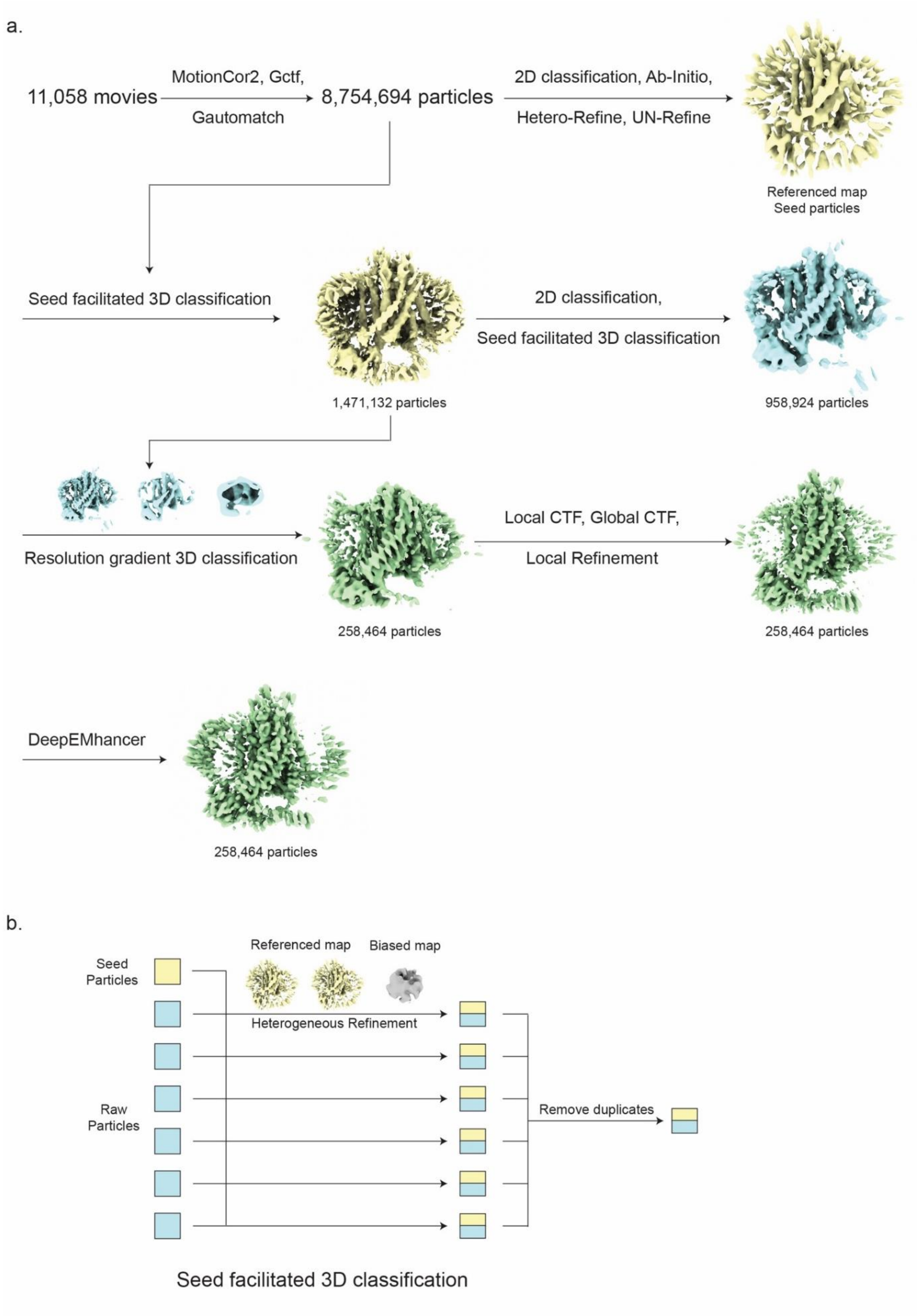


**Supplementary Figure 2 Cryo-EM data processing.** (**a**) Workflow of the TMEM63C data processing. (**b**) Details of seed facilitated 3D classification.

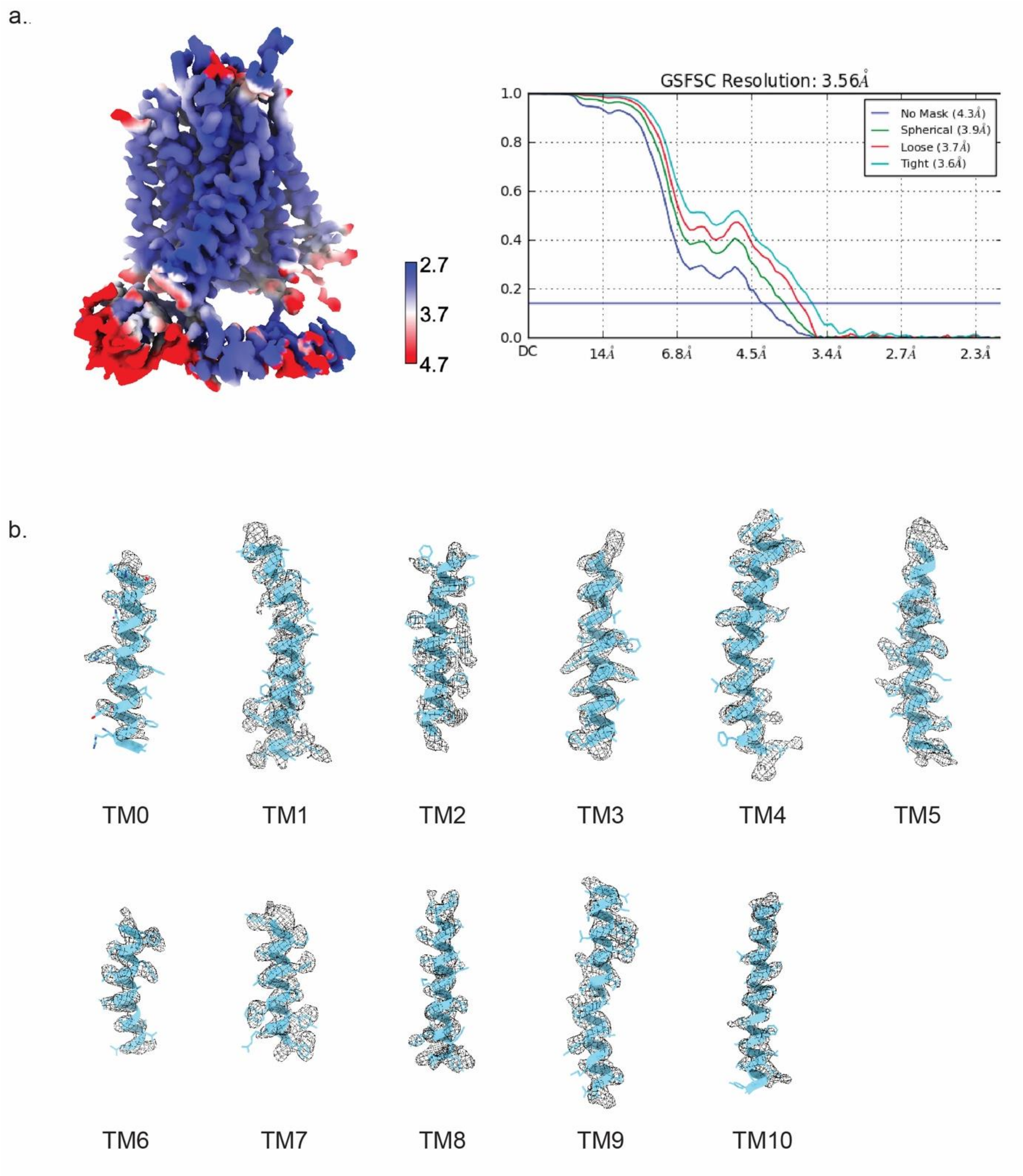


**Supplementary Figure 3 Cryo-EM analysis of TMEM63C.** (**a**) Local resolution estimation of TMEM63C density map (left). FSC curves of the refined model at 3.56 Å resolution (right). (**b**) Cryo-EM density segments (black mesh) and atomic models (blue) of 11 transmembrane α-helices.

| Similarity / Identity | mTMEM63C | mTMEM63B | mTMEM63A | HsTMEM63C | HsTMEM63B | HsTMEM63A | DmTMEM63 | OsOSCA1.2 | AtOSCA1.1 | AtOSCA1.2 | AtOSCA3.1 |
|---|---|---|---|---|---|---|---|---|---|---|---|
| mTMEM63C | 100 | 63 | 60 | 91 | 62 | 63 | 51 | 40 | 39 | 39 | 41 |
| mTMEM63B | 44 | 100 | 75 | 62 | 98 | 75 | 53 | 44 | 41 | 41 | 40 |
| mTMEM63A | 41 | 58 | 100 | 60 | 75 | 95 | 52 | 42 | 40 | 40 | 40 |
| HsTMEM63C | 84 | 42 | 40 | 100 | 62 | 64 | 51 | 42 | 40 | 41 | 39 |
| HsTMEM63B | 43 | 98 | 58 | 43 | 100 | 75 | 54 | 44 | 41 | 42 | 41 |
| HsTMEM63A | 44 | 59 | 90 | 44 | 59 | 100 | 52 | 42 | 39 | 39 | 39 |
| DmTMEM63 | 31 | 33 | 32 | 31 | 33 | 31 | 100 | 41 | 41 | 42 | 39 |
| OsOSCA1.2 | 24 | 25 | 24 | 24 | 25 | 24 | 21 | 100 | 80 | 81 | 53 |
| AtOSCA1.1 | 22 | 21 | 21 | 22 | 22 | 22 | 21 | 69 | 100 | 92 | 52 |
| AtOSCA1.2 | 19 | 21 | 21 | 22 | 21 | 22 | 20 | 69 | 85 | 100 | 51 |
| AtOSCA3.1 | 23 | 24 | 25 | 21 | 24 | 23 | 22 | 33 | 30 | 29 | 100 |

**Supplementary Figure 4** Summary of sequence identity (green) and similarities (blue) of proteins in OSCA/TMEM63 family. mTMEM63C (UniProtKB: Q8CBX0), mTMEM63B (UniProtKB: Q3TWI9), mTMEM63A (UniProtKB: Q91YT8), HsTMEM63C (UniProtKB: Q9P1W3), HsTMEM63B (UniProtKB: Q5T3F8), HsTMEM63A (UniProtKB: O94886), DmTMEM63 (UniProtKB: Q6NP91), OsOSCA1.2 (UniProtKB: Q5TKG1), AtOSCA1.1 (UniProtKB: Q9XEA1), AtOSCA1.2 (UniProtKB: Q5XEZ5) and AtOSCA3.1 (UniProtKB: Q9C8G5).

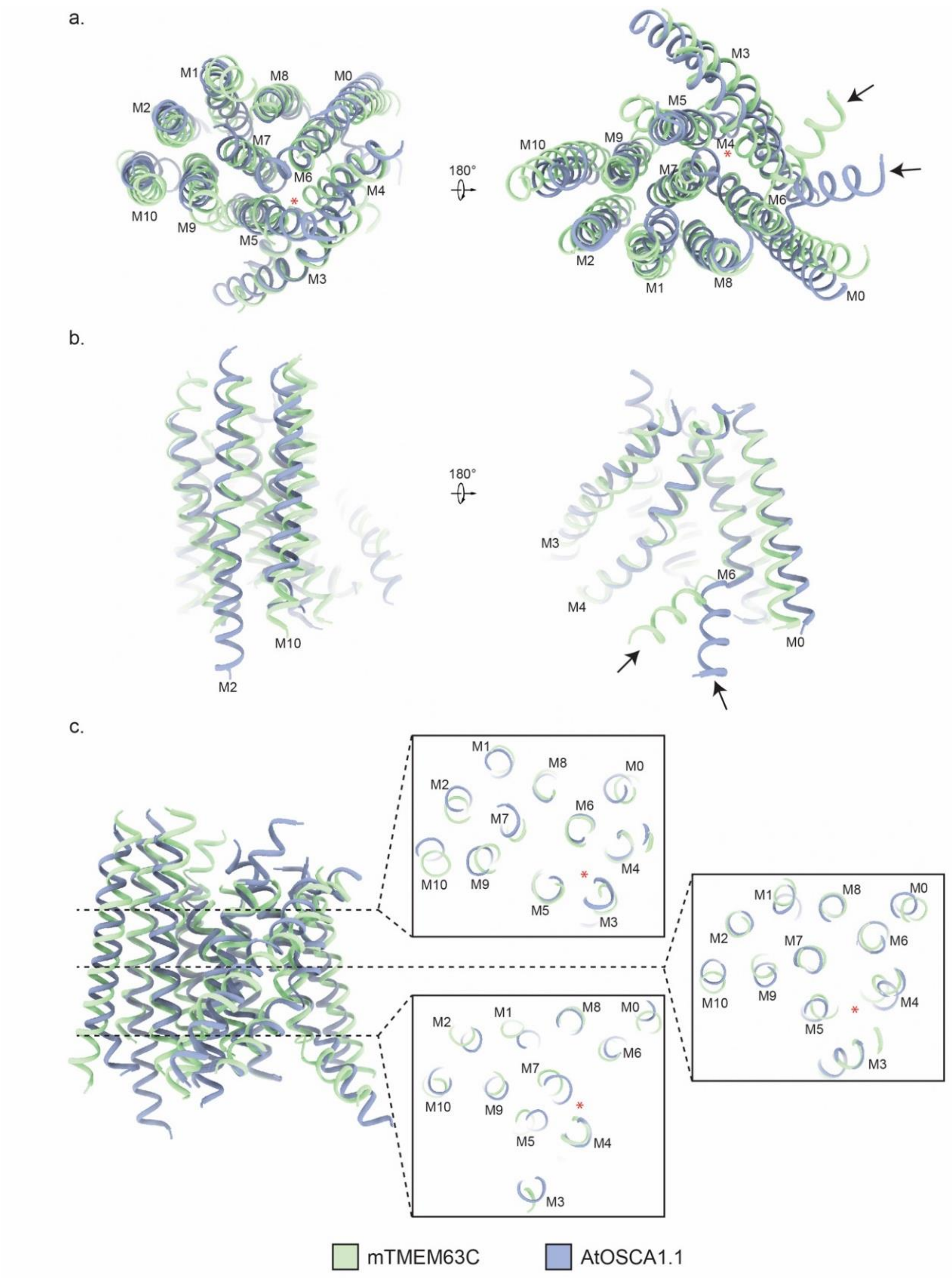


**Supplementary Figure 5 Structure comparison of mTMEM63C and AtOSCA1.1 (PDB:6JPF).** (**a**) Top view (left) and bottom view (right) of mTMEM63C (green) and AtOSCA1.1 (dusty blue). The eleven transmembrane α-helices are labeled with M0 to M10. (**b**) Side views of mTMEM63C and AtOSCA1.1. The M2, M10 and α-helices linked to M6 (black arrows) show conformation changes between mTMEM63C and AtOSCA1.1. (**c**) Cross sections of mTMEM63C and AtOSCA1.1. The red asterisk indicates the pore region.

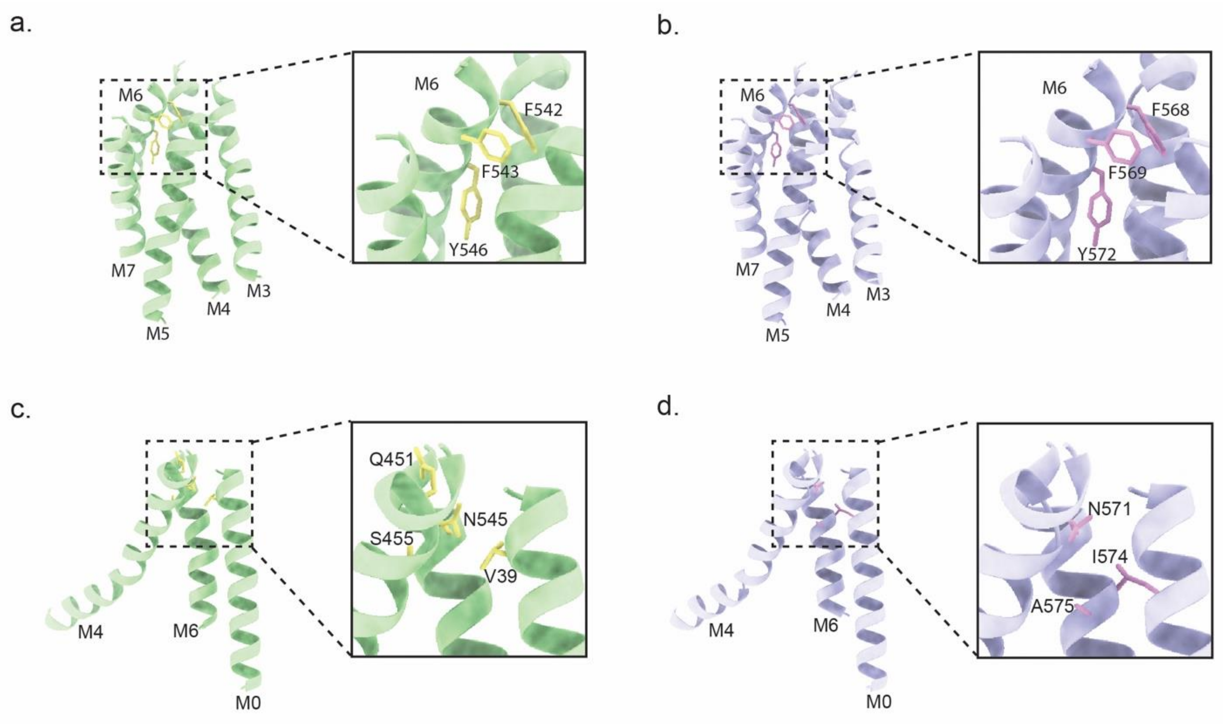


**Supplementary Figure 6 Key residues that are important for gating in TMEM63C and TMEM63B.** (**a**) TMEM63C pore forming α-helices TM3 to TM7 (green). The conserved bulky residues F542, F543 and Y546 (yellow) of TMEM63C in the pore. (**b**) TMEM63B pore forming α-helices M3 to M7 (purple). The conserved bulky residues F568, F569 and Y572 (pink) of TMEM63B in the pore. (**c**) TMEM63C V39 in TM0, Q451, S455 in TM4 and N545 in TM6. (**d**). TMEM63B N571, I574 and A575 in TM6. All the residues are near the extracellular side.

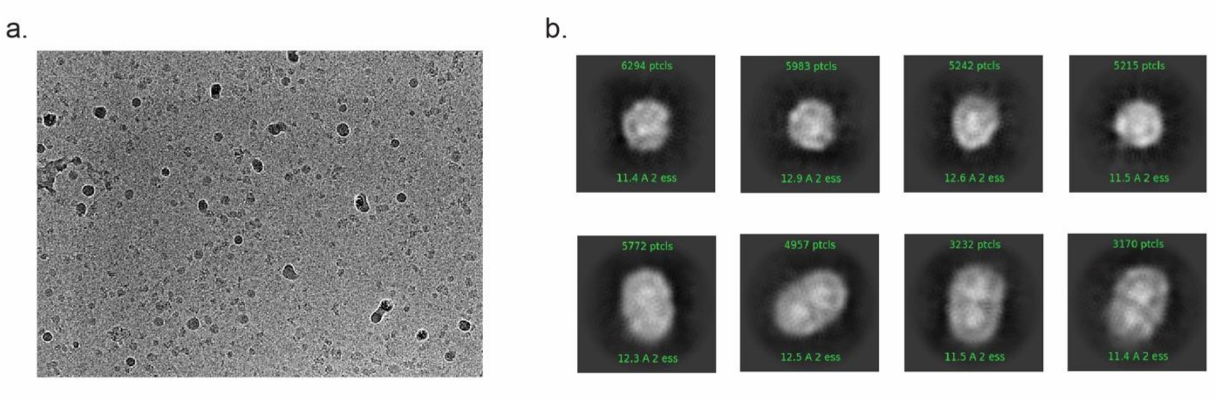


**Supplementary Figure 7 Cryo-EM analysis of TMEM63B.** (**a**) A representative cryo-EM micrograph of TMEM63B. (**b**) The 2D class averages of TMEM63B indicate that both monomer (top) and dimer (bottom) exist.

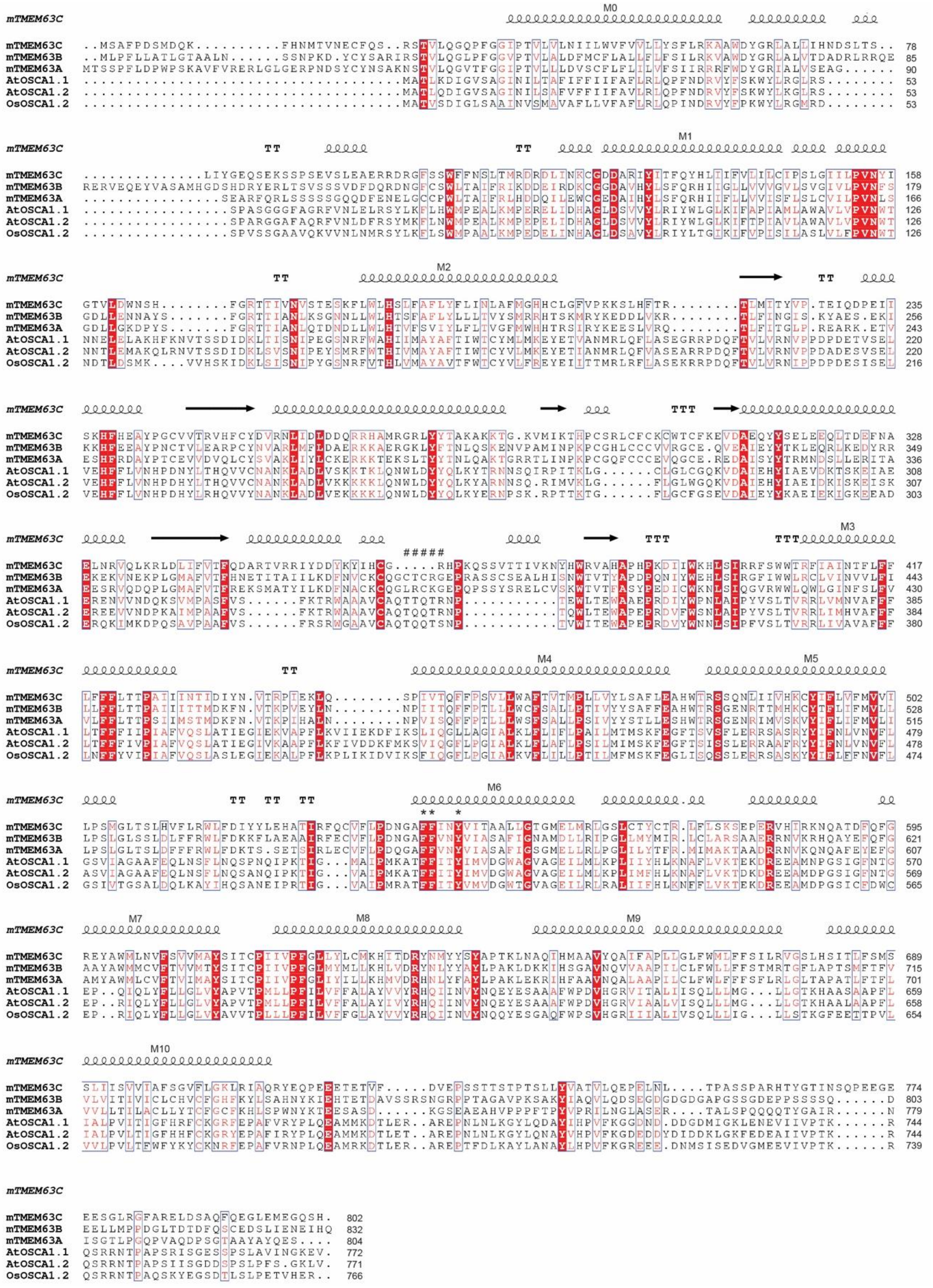


**Supplementary Figure 8 Sequence alignment of mTMEM63C, mTMEM63B, mTMEM63A, AtOSCA1.1, AtOSCA1.2 and OsOSCA1.2.** The conserved residues are colored in red. The secondary structure is shown above the sequences based on the mTMEM63C structure. α-helices are indicated as coil, β-sheets are indicated as arrows, and β-turns are indicated as T. The eleven transmembrane α-helices are labeled with M0 to M10. The symbol “*” shows the three gating residues in the pore, and the symbol “#” shows the OSCA dimer interface residues.

**Supplementary Table 1 Cryo-EM data collection and model building**

| | |
|---|---|
| **Data collection** | |
| EM equipment | Titan Krios |
| Voltage (kV) | 300 |
| Detector | Gatan K3 summit |
| Magnification | 81,000 x |
| Pixel size (Å) | 1.06 |
| Electron dose (e/Å$^2$) | 50 |
| Defocus range (μm) | 1.0~2.5 |
| Collected movies | 11,058 |
| **Reconstruction** | |
| Software | cryoSPARC, RELION |
| Final particles | 258,464 |
| B-factors (Å$^2$ ) | -113.5 |
| FSC threshold | 0.143 |
| Map resolution (Å) | 3.56 |
| EMDB | EMD-XXXX |
| PDB | XXXX |
| **Validation** | |
| RMSD length (Å) | 0.004 |
| RMSD Angles (°) | 0.865 |
| Clash score | 14.96 |
| Outliers | 0 |
| Allowed | 8.01 |
| Favored | 91.99 |
